# Revealing hidden knowledge in amnestic mice

**DOI:** 10.1101/2025.01.09.632026

**Authors:** Andrea Santi, Sharlen Moore, Kelly A. Fogelson, Aaron Wang, Jennifer Lawlor, Jordan Amato, Kali Burke, Amanda M. Lauer, Kishore V. Kuchibhotla

## Abstract

Alzheimer’s disease (AD) is a form of dementia in which memory and cognitive decline is thought to arise from underlying neurodegeneration. These cognitive impairments, however, are transient when they first appear and can fluctuate across disease progression. Here, we investigate the neural mechanisms underlying fluctuations of performance in amnestic mice. We trained APP/PS1+ mice on an auditory go/no-go task that dissociated learning of task contingencies (knowledge) from its more variable expression under reinforcement (performance). APP/PS1+ exhibited significant performance deficits compared to control mice. Using large-scale two-photon imaging of 6,216 excitatory neurons in 8 mice, we found that auditory cortical networks were more suppressed, less selective to the sensory cues, and exhibited aberrant higher-order encoding of reward prediction compared to control mice. A small sub-population of neurons, however, displayed the opposite phenotype, reflecting a potential compensatory mechanism. Volumetric analysis demonstrated that deficits were concentrated near Aβ plaques. Strikingly, we found that these cortical deficits were reversed almost instantaneously on probe (non-reinforced) trials when APP/PS1+ performed as well as control mice, providing neural evidence for intact stimulus-action knowledge despite variable ongoing performance. A biologically-plausible reinforcement learning model recapitulated these results and showed that synaptic weights from sensory-to-decision neurons were preserved (i.e. intact stimulus-action knowledge) despite poor performance that was due to inadequate contextual scaling (i.e. impaired performance). Our results suggest that the amnestic phenotype is transient, contextual, and endogenously reversible, with the underlying neural circuits retaining the underlying stimulus-action associations. Thus, memory deficits commonly observed in amnestic mouse models, and potentially at early stages of dementia in humans, relate more to contextual drivers of performance rather than degeneration of the underlying memory traces.

## Main

Over 55 million people worldwide suffer from Alzheimer’s disease (AD), the most common form of dementia (WHO, 2022). Before the start of large-scale neurodegeneration, the progressive accumulation of amyloid-beta (Aβ) peptides and tau tangles, together with other pathological changes, impact the functional integrity of neural circuits leading to cognitive and memory deficits that worsen with time^1^. These cognitive impairments, however, are transient when they first appear, and become longer, more frequent, and eventually seemingly permanent at later stages of the disease. Interestingly, even at these later stages, increasing evidence shows that patients with AD and related dementias also experience positive cognitive fluctuations^2,3^, where memories and cognitive abilities temporarily improve—including the paradigmatic case of lucid intervals^4–7^. Despite the abundant body of work exploring synaptic and molecular changes associated with cognitive deficits, less is known about the circuit-level mechanisms underlying these impairments either in patients or amnestic animal models, and whether these alterations are permanent, or context-dependent, as some clinical observations suggest.

Cognitive performance can be temporarily impacted by factors such as stress, anxiety or agitation, comorbid symptoms that are commonly observed in AD patients^8–11^ as well as animal models of AD^12–18^, since cognitive performance itself, is highly sensitive to external context and internal state^19–23^. Recent work has shown that knowledge of a task and performance are not the same, such that animals may have latent knowledge of stimulus-action associations—revealed on non-reinforced trials—that is obscured by traditional performance measures under reinforcement^22,24–26^. This builds on a long-known (though often overlooked) phenomenon in learning theory which is sometimes referred to as an ‘extinction burst’, where there is a temporary increase in the ‘strength’ of a behavior when an expected reinforcer is removed^27–30^. Performance is thus a function of both knowledge of a task—the underlying strength of the stimulus-action associations—and non-associative contextual factors.

Here, we leverage this behavioral manipulation to investigate the neural mechanisms underlying fluctuations of performance in amnestic mice. We focused neural interrogations on the auditory cortex, as it is the first site in the auditory system with amyloid accumulation in these mice ^31^, plays a key role in enhancing the detection of discrimination of auditory cues, and integrates higher-order behavioral signals to enable learning and performance^32–39^. Moreover, there is increasing evidence linking central auditory deficits and dementia^40–48^, even before the manifestation of cognitive decline^43^.

## Results

### Excessive suppression and compensatory facilitation of auditory cortical neurons in amnestic mice performing a go/no-go task

To explore the impact of amyloid accumulation on the neural computations underlying cognitive performance, we trained 6 to 8-month-old APP/PS1+ on an auditory go/no-go task (Fig. **1a-b**). These mice show significant amyloid accumulation due to the overexpression of amyloid precursor protein (APP) in combination with mutant presenilin 1 (PS1;^31,49–52^). Head-fixed mice learn to lick to a target tone (S+) to obtain a water reward and withhold from licking to a different tone (S-) to avoid a time-out. In line with the results obtained from other cognitive tests in amyloid models^10,53,54^, adult APP/PS1+ mice (n=13) exhibited significant performance deficits in this task when compared to age-matched controls (APP/PS1-, n=12, Fig. **1c**). We then performed large-scale *in vivo* two-photon calcium imaging with single-neuron resolution in a subset of these mice after they reached a final stable performance (Fig. **1a, d**). We imaged 6,216 neurons from 8 mice (n=5 APP/PS1+, n=3 control) and found that neurons from APP/PS1+ mice had an overall decrease in responsivity (53.85% APP/PS1+; 76.9% control, χ2=317.5019, p<0.0001, Fig. **1e**). We next analyzed the stimulus selectivity of these neurons, as some neurons exhibited higher evoked responses to the S+ and others to the S- (S+ selective vs. S-selective; Fig. **1f**). We observed an overall decrease in stimulus selectivity for all neurons (Fig. **1g**, *left*) in APP/PS1+ mice. This decrease was true for both S+ and S-selective neurons (Fig. **1g-h**) and was driven by a higher percentage of tone-suppressed neurons in APP/PS1+ mice that were significantly less selective (1279 out of 2177 -58.8%- vs 322 out of 1671 -19.3%-of tone-suppressed cells in APP/PS1+ vs controls; Fig. **1h-i**). Interestingly, while the percentage of activated neurons was significantly reduced (by almost half) in the APP/PS1+ mice (41.2 vs 80.7% of tone-responsive cells in APP/PS1+ vs controls; χ2=604.8955, p<0.0001; Fig. **1h**), the smaller percentage of activated neurons showed higher stimulus selectivity (Fig. **1i**, *bottom*) in APP/PS1+ mice, and increased stimulus-evoked responses (S+, 27.2% higher in APP/PS1+; S-, 65.1% higher in APP/PS1+), even when compared to neurons from control mice. These results suggest that the cortical network ‘compensates’ for the emerging deficits by enhancing the activity and selectivity of the remaining activated population.

**Figure 1.**
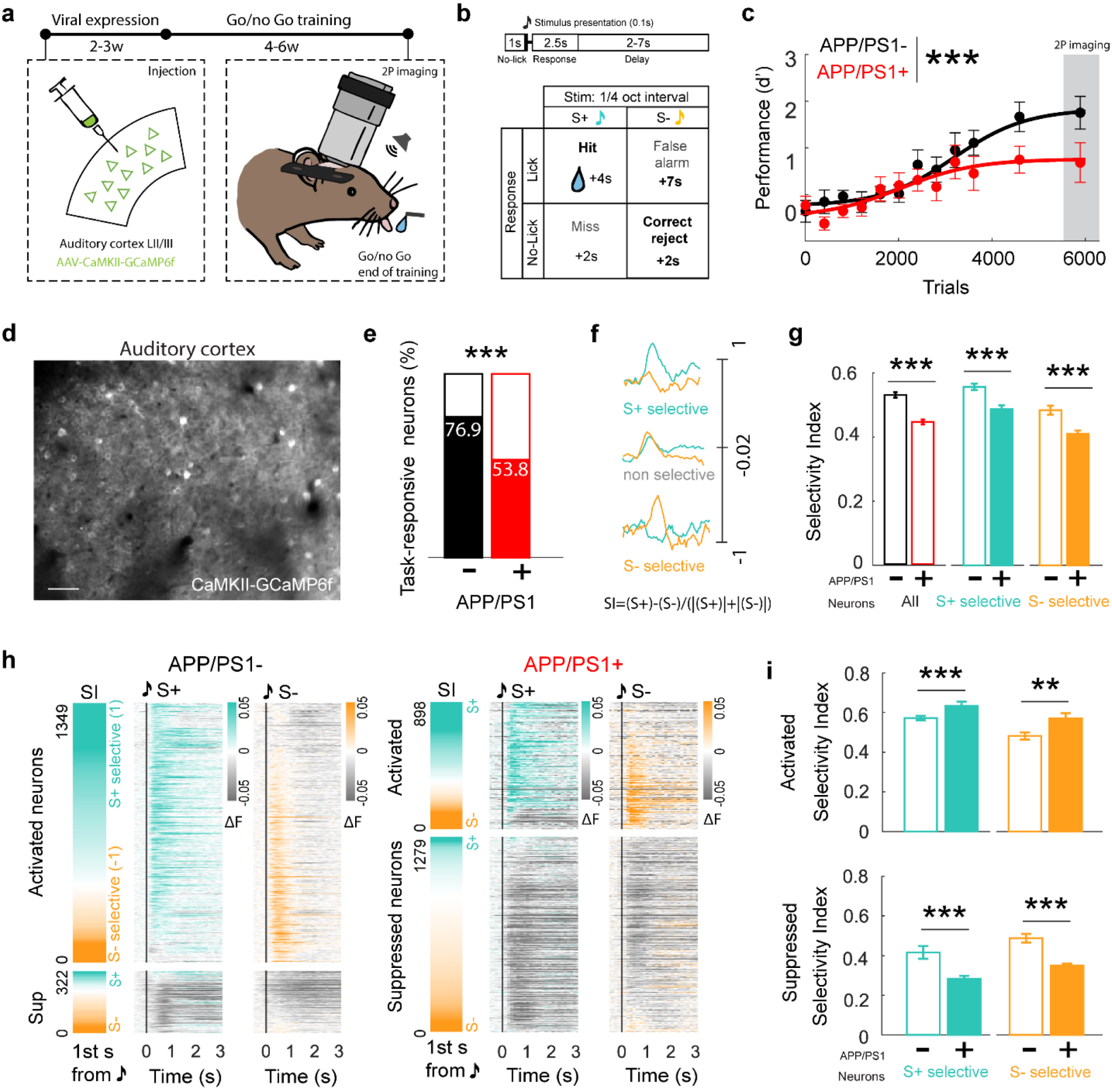
Excessive suppression and compensatory facilitation in auditory cortical neurons of 6-8mo APP/PS1+ mice with performance deficits. a, Schematics of experimental approach. b, Trial structure and possible outcomes. c, APP/PS1+ mice (red) show significantly impaired performance over time in the auditory go/no go task (d’=z(false alarms/S-trials) – z(hits/S+trials); Mixed-effects model; TimexGenotype, F(59, 1300)=1.960, p<0.0001). d, Representative image of recorded field of view of excitatory neurons in the auditory cortex. e, APP/PS1+ mice exhibit fewer tone-responsive neurons (APP/PS1-1671/2173 vs APP/PS1+ 2177/4043; χ2=317.5019, p<0.0001). f, Examples of stimulus selective neurons. g, Stimulus selectivity of excitatory neurons is reduced overall (red bar; Z=8.1777, p=3.4939e+06) and in S+ and S-selective neurons (Selectivity in S- selective neurons displayed in absolute levels; S+, n=1047 APP/PS1- and n=964 APP/PS1+, Z=4.8816, p=1.0525e-06; S-, n=624 APP/PS1- and n=1213 APP/PS1+, Z=-4.4925, p=7.0390e-06, Wilcoxon rank sum test). h, Heatmap of significantly responsive neurons in response to the S+ and the S- tones in control (*left*) and APP/PS1+ (*right*) mice, sorted by selectivity index (first column; calculated over the first second after tone presentation). i, *Top*, increase in selectivity of the few activated neurons compared to control mice (S+, n=942 APP/PS1- and n=562 APP/PS1+, Z=-3.5277, p=4.1912e-04; S-, n=407 APP/PS1- and n=336 APP/PS1+, Z=2.7133, p=0.0067, Wilcoxon rank sum test). *Bottom*, suppressed neurons in APP/PS1+ mice exhibit far less selectivity than suppressed neurons in control mice (S+, n=105 APP/PS1- and n=402 APP/PS1+, Z=3.9667, p=7.2866e-05, S-, n=217 APP/PS1- and n=877 APP/PS1+, Z=-6.0897, p=1.1309e-09, Wilcoxon rank sum test). ∗ p<0.05, ∗∗ p<0.01, ∗∗∗p<0.001, ns=non-significant.

One possible explanation for these deficits is that amyloid impacts feedforward sensory processing, independent of the task. To test this, we measured sound-evoked responses and selectivity before any task training. We found only a modest decrease (∼5%) of tone-responsive neurons pre-training (Extended Data Fig. 1a), while stimulus selectivity was higher compared to controls in the passive context and APP/PS1+ mice during the task (Extended Data Fig. 1c). Additionally, the majority of responsive neurons of APP/PS1+ mice were activated (Extended Data Fig.**1b**; 79.6% in the passive context vs. 41.2% during the task), suggesting the observed neural effects during behavior are not resulting from deficits in sensory processing. To further test this, we measured auditory brainstem responses, a standard electrophysiological approach to assess potential peripheral or subcortical hearing impairments^55^. Studies have demonstrated conflicting results about the extent to which various AD models exhibit peripheral forms of hearing loss^56–59^. We found that detection thresholds were not different between APP/PS1+ and control mice for all the frequencies selected to train the mice and were below the behavioral stimulus sound levels (<65dB SPL) (Extended Data Fig. 2a-b). Moreover, consistent with the lack of subcortical amyloid accumulation in this model^31^, we found no significant changes in the latencies or amplitudes of the ABR peaks for any stimulus at 70dB SPL (Extended Data Fig. 2c-d; Supplementary Table **1**).

We then used a linear decoder to test whether these neural alterations impacted population decoding of the two stimuli. We found that stimulus decoding was impaired immediately after tone-onset (‘early-in-trial’) and worsened late-in-trial (Extended Data Fig. 3a). Importantly, we observed a reduced number of neurons that contributed to the stimulus decoding in APP/PS1+ mice, indicating that behavioral encoding becomes less distributed and more concentrated in cortical networks with significant amyloid accumulation (Extended Data Fig. 3a-b). Together, these results point to excessive suppression (decreasing overall responsivity and selectivity) and a surprising level of compensatory facilitation that counteracts and partially preserves population-level stimulus-related decoding (Extended Data Fig. 3a).

### Aberrant reward prediction activity in auditory cortical neurons of amnestic mice

Detailed inspection of the neural activity traces showed that neurons in the auditory cortex not only exhibited classical stimulus-evoked responses (‘early-in-trial’ activity) but also exhibited prolonged ‘late-in-trial’ activity (Fig. **2a**). In addition, population decoding was significantly impaired late-in-trial (Extended Data Fig. 2a), prompting us to explore whether late-in-trial activity patterns could be encoding other task-relevant information^26,32,60–64^. We analyzed the peak activity of every neuron in hit trials (correct behavioral response to S+) and we found that, while most neurons had only one peak of tone-evoked activity early-in-trial, some exhibited a peak of activity late-in-trial (Fig. **2a-b**). Similar to the stimulus-evoked activity patterns, the APP/PS1+ network showed fewer late-in-trial activated neurons in response to the S+ tone (6.9%, 280 out of 4,043 neurons) compared to neurons from control mice (14.5%, 315 out of 2,173; χ2=25.4865 test; p<0.0001; Fig. **2c**).

**Figure 2.**
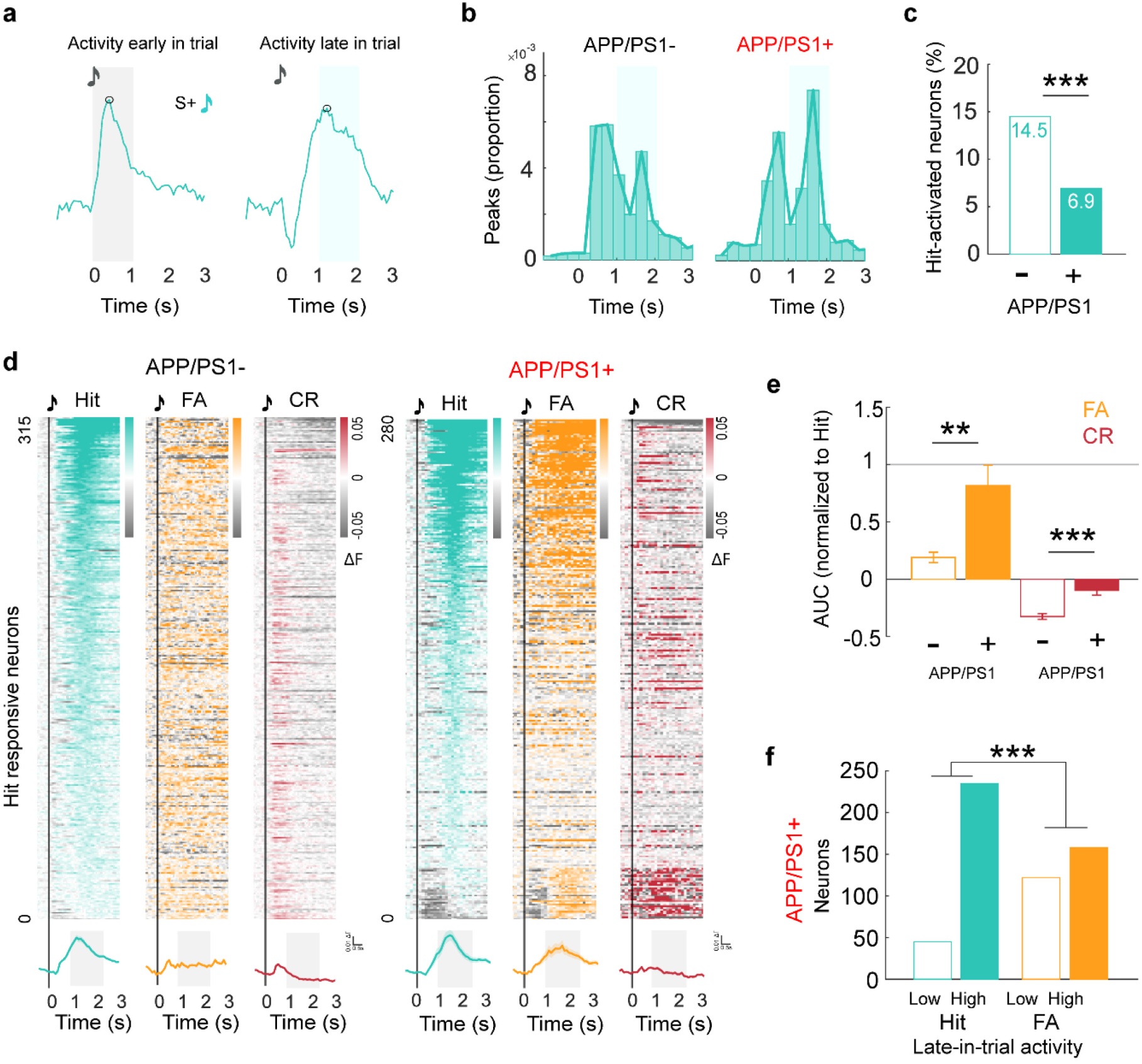
Aberrant reward prediction activity on correct and error trials of 6-8mo APP/PS1+ mice. **a,** Neural responses after tone presentation show different peaks of activity. Black circles indicate detected peaks. Light grey shading indicates tone-evoked activity early-in-trial. Light blue shading indicates behavioral-related late-in-trial activity. **b,** Distribution of peak activity in cells with late**-**in**-**trial significant responses. **c,** Percent of activated neurons late-in-trial in hit trials is significantly decrease in APP/PS1+ network (APP/PS1- 315/2173 vs APP/PS1+ 280/4043; χ2=92.7105; p<0.0001). **d,** *Top*, heatmap of responses by trial type of hit- responsive neurons sorted by late-in-trial activity in hit trials. *Bottom*, averaged neural activity to different trial types of neurons with late-in-trial activity on hit trials in control and APP/PS1+ mice. Gray shading indicates time use to calculate AUC in E. **e,** APP/PS1+ neurons with late-in-trial activity show aberrant activity in false alarm and correct reject trials compared to control neurons (Normalized to hit trials. FA p=0.007; CR p<0.0001, Wilcoxon rank sum test). **f,** 44% of APP/PS1+ neurons show low reward prediction activity on incorrect trials (False alarm; 158 out of 280; χ2=49.2840, p=2.2146e-12). ∗ p<0.05, ∗∗ p<0.01, ∗∗∗ p<0.001, ns=non-significant.

Interestingly, this late-in-trial activity was present on rewarded trials (hits) but not during correct rejections (correct response inhibition during S- trials; **2d-e**) or miss trials (Extended Data Fig. 4), and was independent of lick vigor (Extended Data Fig. 5), suggesting this neuronal ensemble could be encoding reward prediction (see also^26^ which demonstrates that this activity is not driven by licking, licking initiation, or reward consumption). Surprisingly, the late-in-trial responsive neurons of APP/PS1+ mice exhibited an overall higher activity on action-related errors (false alarms, licks in response to the S-), and significantly less suppression in correct rejection trials (Fig. **2d-e**). These analyses indicate that excitatory neurons in amnestic mice display aberrant reward prediction activity for correct and error trials during performance under reinforcement. However, while the higher activity during false alarms in this ensemble of APP/PS1+ neurons was present in the majority of cells, a few of them also showed low activity (Fig. **2f**), suggesting that the network still reflected discriminative encoding of the stimulus-action and stimulus-outcome associations, despite the aberrant reward prediction activity.

### Neural deficits are concentrated near amyloid plaques

Fibrillar and soluble forms of Aβ are highly enriched in the plaque periphery and have been shown to be particularly damaging to neural structure and function^65–70^. We next sought to determine the extent to which the neural deficits we observed were broadly distributed or, instead, concentrated near Aβ plaques. To do this, we injected an amyloid-binding fluorescent dye (Methoxy-X04) within 24 hours of each imaging session^71^. This allowed us to visualize amyloid plaques throughout the field of view (Fig. **3a**). We then performed structural imaging of a 3-dimensional volume to precisely measure the minimum distance of each neuron to the closest plaque. Importantly, our functional imaging site was centered within this volume and the 3-D structural imaging of Aβ plaques was done in a ‘zoomed out’ approach, ensuring an accurate measure of the 3-D distance even for neurons at the edge of the functional imaging site (Fig. **3a**). This analysis revealed a tight correlation between neural responsivity and distance to plaque such that neurons closest to plaques exhibited the weakest stimulus-evoked and late-in-trial signaling (Fig. **3b-c**). Despite this decrease in overall responsivity, the small population of ‘activated’ neurons (those exhibiting a stimulus-evoked increase early-in-trial) remained functionally intact near plaques and exhibited a large compensatory increase in selectivity further from plaques (Fig. **3d**). The larger population of ‘suppressed’ neurons exhibited reduced selectivity near plaques, but intact selectivity further away (Fig. **3e**). These data further suggest that neural dysfunction is concentrated near amyloid plaques while also pointing to novel compensatory processes.

**Figure 3.**
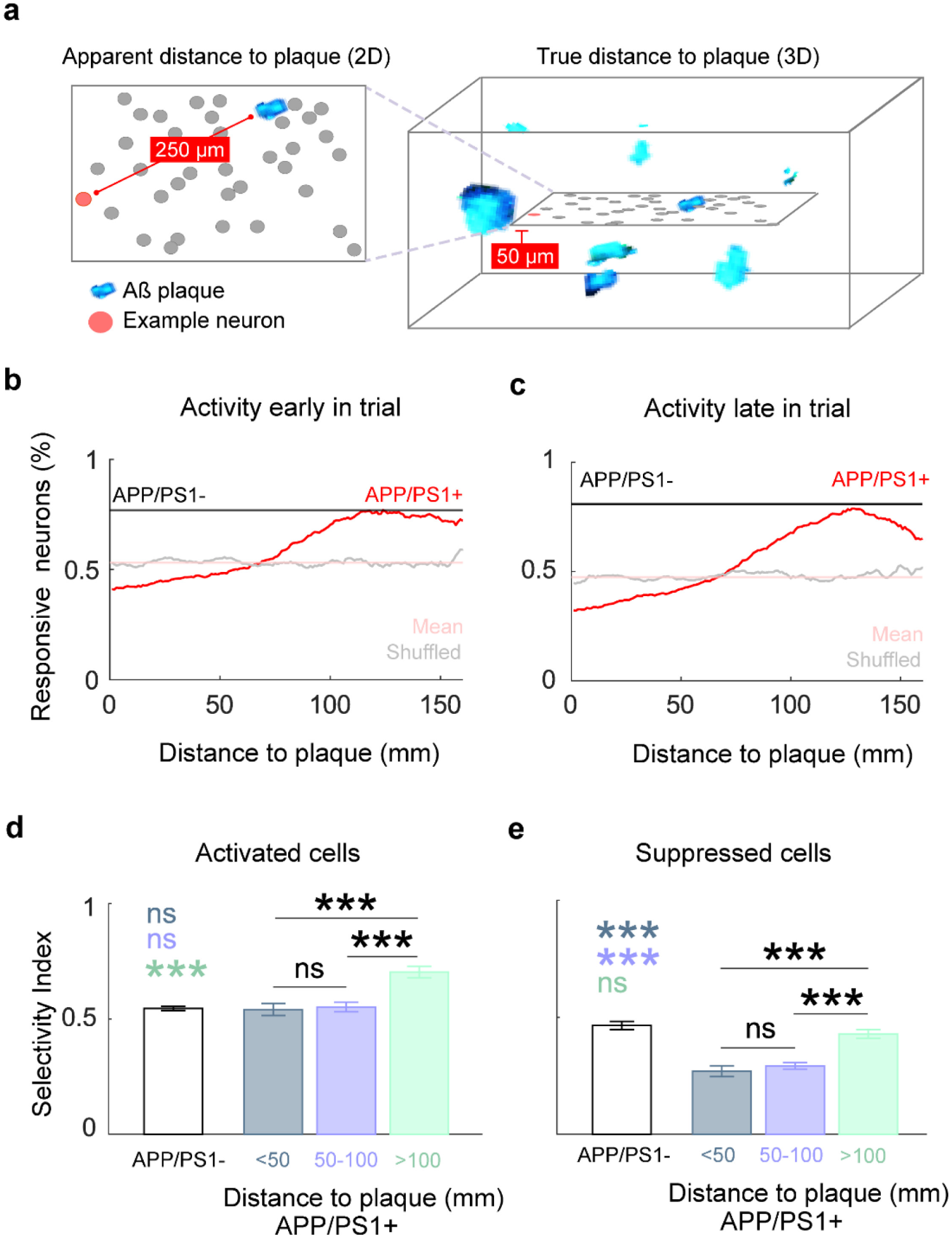
Neural encoding in the auditory cortex of 6-8mo amnestic mice is affected by amyloid plaque proximity. **a,** Schematics of distance to plaque quantification. Plaques in the z-stack light blue. Neuronal ROI in grey **b-c,** Neural responsiveness early (**b**) and late-in-trial (**c**) increases far from plaques. **d,** Selectivity index increases with distance to plaque among activated neurons (APP/PS1- (n=1349) vs APP/PS1+ <50μm (n=205), p=0.6236, vs APP/PS1+ 50-100μm (n=303), p=0.7894, vs APP/PS1+ >100μm (n=188), p=0.2067e-08; Wilcoxon rank sum test. Kruskal-Wallis all APP/PS1+ groups; χ2=25.5918, p=2.7721e-06) and suppressed neurons (**e**) (APP/PS1- (n=322) vs APP/PS1+ <50μm (n=151), p=2.1697e- 11, vs APP/PS1+ 50-100μm (n=410), p=2.3173e-14, vs APP/PS1+ >100μm (n=304), p=0.1189; Wilcoxon rank sum test. Kruskal-Wallis all APP/PS1+ groups; χ2=45.0647, p=1.6381e-10). ∗ p<0.05, ∗∗ p<0.01, ∗∗∗p<0.001, ns=non-significant.

### Neural deficits and behavioral performance are restored in non-reinforced trials

Cognitive performance in a task depends on both knowledge of the task (the underlying associations) and non-associative contextual factors. Here, we tested whether performance deficits in amnestic mice were driven by one, the other, or both. We exploited non-reinforced probe trials to better assess the strength of the underlying stimulus-action associations. We reasoned that if knowledge of the task was impaired, performance on these probe trials would be low (just like on reinforced trials). Alternatively, if task knowledge is intact but non-associative contextual factors were impaired, performance on probe trials would be significantly higher than on reinforced trials. To do this, we interleaved short blocks of non-reinforced trials throughout learning and plateau performance in amnestic and control mice (Fig. **4a**). Surprisingly, we found that performance on probe trials was strikingly higher than on reinforced trials (Fig. **4b**), and similar to the performance of control mice (Supplementary Table **2)**.

**Figure 4.**
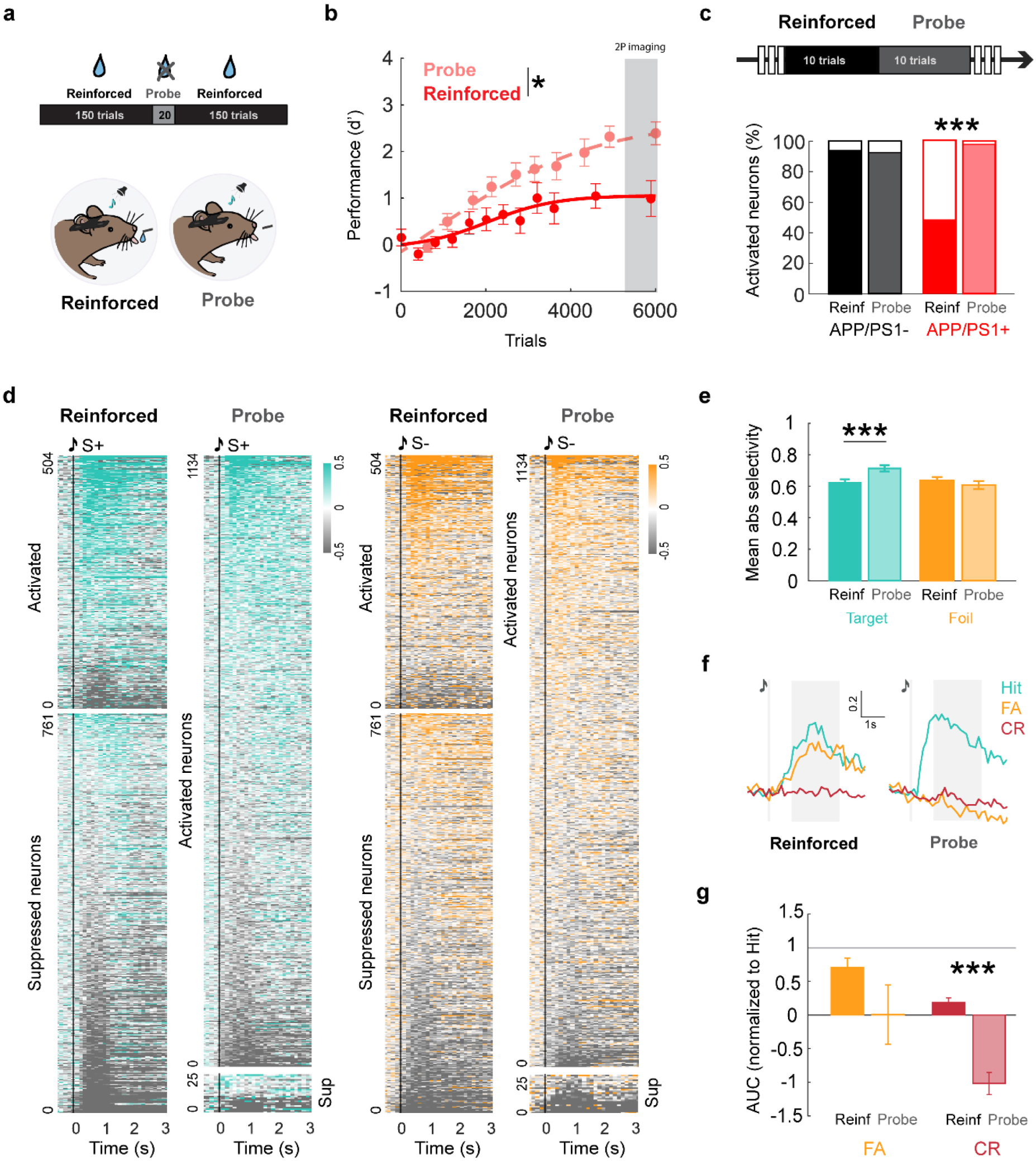
Neural dynamics of 6-8mo APP/PS1+ are restored in the probe context. **a,** Every other day, a subset of 20 trials were administered in the middle of the training session were all task settings were maintained but reward was omitted (‘probe trials’). **b,** APP/PS1+ mice performed significantly better in task blocks were reinforcement was not delivered (Probe context; TimeXContext, F(9,207)=2.104, p=0.0305; Time, F(2.098,48.25)=19.07, p<0.0001; Context, F(1,24)=5.242, p=0.0311). **c,** *Top*, 10 reinforced trials immediately before probe were selected to compare responsiveness to S+ or S-. Bottom, percent of activated neurons in probe is increased (filled portion of bar) while suppressed cells (empty portion) are decreased in APP/PS1+ (Reinforced vs Probe χ2=925.9836, p<0.0001). Distribution of activated and suppressed neurons in APP/PS1- mice is maintained (Reinforced vs Probe; χ2=0.2321, p=0.630). **d,** Heatmap of significantly responsive neurons in reinforced and probe contexts sorted by activity. **e,** Tone-responsive neurons increase their mean selectivity in probe context (S+, p=0.0005; S-, p=0.3762, Wilcoxon rank sum test). **f,** Example APP/PS1+ neuron with putative reward prediction activity show increased suppression in S- trials in probe compared to reinforced context. **g,** Quantification of late-in-trial activity for all hit- significantly-responsive APP/PS1+ neurons (Normalized to hit trials. FA p=0.1367, CR p<0.0001). ∗ p<0.05, ∗∗ p<0.01, ∗∗∗ p<0.001, ns=non-significant.

These data suggest that performance deficits in amnestic mice at this age (6-8mo) are not related to the strength of the underlying associations (i.e. task knowledge). We next sought to assess whether the performance deficits under reinforcement could arise from disengagement, impulsivity, or impaired motor abilities. We tested each of these systematically and found no evidence of disengagement (Extended Data Fig. 6a, no difference in hit rates), impulsivity (Extended Data Fig. 6b, no difference in reaction times) or motor-related licking ability (Extended Data Fig. 6a-b). Altogether, these results point to deficits in contextual integration rather than degeneration of the underlying associations.

Given that 6-8mo APP/PS1+ mice exhibited evidence of intact associations on probe trials, we next investigated neural activity in the auditory cortex. Remarkably, we found that the proportion of activated APP/PS1+ neurons was significantly higher on probe trials compared to reinforced trials, while the proportion of significantly suppressed neurons was strongly reduced (Fig. **4d**, *bottom*). Moreover, the APP/PS1+ network was more selective to the S+ tone in the probe context when compared with the selectivity of the same cells in the reinforced context (Fig. **4e-f**). We then analyzed reward prediction neurons from APP/PS1+ mice (i.e. neurons with activity late-in-trial on hit trials but also incorrectly on false alarm trials) and observed a reduced reward prediction activity on error trials (i.e. false alarms), and increased suppression on correct reject trials (Fig. **4g**) resembling the activity of control mice (Fig. **1g-i**). Finally, we observed that mice could sometimes perform better on the task on reinforced trials over shorter time-scales, i.e. 100-trial blocks. To test whether this improved performance was driven by the same putative mechanism— restored reward prediction activity—we correlated behavioral performance blocked into 100-trial bins with stimulus decoding from the ‘late-in-trial’ period. We found a strong correlation between the two (Extended Data Fig. 3d)., suggesting that transient periods of higher performance are not random and are potentially driven by improved cortical function. These data suggest that network deficits (reduced responsivity and selectivity, aberrant reward prediction activity) can be reversed almost instantaneously in the probe context (and to some extent on blocks of reinforced trials), providing neural evidence for the persistent strength of the underlying associations.

To better understand the computational basis of these contextual deficits in performance, we then applied a biologically plausible reinforcement learning model to behavioral data from the APP/PS1+ and control mice of 6-8mo^22,72^ (Extended Data Fig. 7a). In this model, sensory neurons (S+, S-, S) project to a read-out population consisting of a decision neuron (D) and a modulatory inhibitory neuron (I). This model identifies associative strength (task knowledge) as the synaptic weights between sensory-to-decision neurons while capturing contextual variability in performance via a single contextual scaling parameter that is only applied to the decision read- out (without impacting the underlying synaptic weights). The model accurately recapitulated reinforced and probe performance for APP/PS1+ and control mice (Extended Data Fig. 7b-c). By running several model simulations over each individual animal, we obtained the distributions of each model parameter that best capture individual animal behavior. In line with our behavioral and neural data, we found that the behavioral deficits in 6-8mo APP/PS1+ mice were largely explained by changes in contextual and synaptic weight scaling and the corresponding inhibitory weights (Extended Data Fig. 7d). Other parameters, such as excitatory weights or the S+ or S- learning rate remained unchanged in these mice at early stages of the disease (Extended Data Fig. 7d). These results indicate that 6-8mo APP/PS1+ mice exhibit no degradation of the synaptic weights underlying the associative knowledge.

Finally, we performed similar behavioral studies in 2-3mo (young) and 10-12mo (aging) mice to identify how amyloid affects contextual performance through aging. We found that both age and genotype impacted performance (Extended Data Fig. 8), since both control and APP/PS1+ mice showed an age-related impairment in the reinforced contexts (Extended Data Fig. 8a,c). Interestingly, while control mice exhibited little to no age-related decline on probe trials, APP/PS1+ mice, exhibited an age-related decrease in performance on both reinforced and probe trials (Extended Data Fig. 8b,d). These results were recapitulated by our reinforcement learning model that showed that in addition to deficits in the contextual scaling parameter, learning rates and excitatory weights also became impacted at 10-12mo in APP/PS1+ (Extended Data Fig. 7e). Taken together, these data suggest that performance deficits that worsen during aging relate to aberrant integration of non-associative contextual factors, and that this is accelerated by increasing levels of amyloid and take place before the weakening of the underlying associations.

## Discussion

Here, we exploit non-reinforced probe trials to demonstrate that APP/PS1^+^ mice performing an auditory go/no-go task exhibit profound performance deficits on reinforced trials but completely intact task knowledge (probe trials) even after significant amyloid deposition (6-8-months-old). Importantly, interrogation of neural dynamics in the auditory cortex of APP/PS1+ mice demonstrated that these performance deficits are driven by an increased suppression of neural activity (reduced overall responsivity and higher percentage of suppressed neurons), reduced stimulus selectivity, and aberrant higher-order encoding of reward prediction. These neural deficits were endogenously and transiently restored on probe trials and during short blocks of higher performance during reinforcement.

These results support the idea that prodromal cognitive impairments are due to retrieval deficits^73,74^ and that the memory trace is present but silent (or suppressed) when amnestic mice show these impairments^74,75^. Strikingly, our results suggest that the silencing of the memory trace (1) is not permanent, (2) can be endogenously reversed nearly instantaneously by changing the behavioral context, and (3) that recovery is neurally instantiated as restored responsiveness, stimulus selectivity, and behavioral integration even in the presence of substantial fibrillar and soluble amyloid. This provides some of the first evidence that apparently silent memory traces can be re-engaged without exogenous perturbations, and points to restoration of sensory and higher-order neural encoding as drivers of transient improvements in cognition.

Interestingly, our data also show evidence for neural compensation (in the form of a sub- population of neurons exhibiting enhanced activity and stimulus selectivity). These context-dependent effects might reflect alterations in the integration of ascending cholinergic neuromodulation (recruited in highly motivated states such as the reinforced context)^76–79^ and their interaction with inhibitory micro-circuits in the cortex^80,81^. In particular, ascending neuromodulation recruits different interneurons that suppress task-irrelevant excitatory neurons while amplifying task-relevant excitatory neurons in the cortex^82^. One area of future investigation will be to test whether amyloid disrupts the integration of cholinergic inputs, disturbing the delicate balance between inhibition (via PV+ interneurons^83^) and disinhibition (via VIP+ interneurons) and thereby impairing the activation of the memory trace^84^.

It is important to note that APP/PS1+ mice do not provide a complete model of AD. These mice do not exhibit large-scale neurodegeneration nor intracellular tau tangles; in addition, these (and other) transgenic lines have overexpression artifacts. With that said, our most striking findings relate not to a deficit, but to the transient and contextually-triggered access to surprisingly intact task knowledge. Moreover, these mice remain valuable models of amyloidosis and prodromal, and even preclinical, AD, especially when there is convincing evidence of age- and amyloid-dependent phenotypes, which we observe. It will be important for future studies to extend our approach to additional models and translate these findings to behavioral and neurological testing in humans. Finally, although further work is required to elucidate the precise mechanisms leading to the observed deficits and compensatory processes, our reinforcement learning model supports the role of contextual scaling as the main driver of poor cognitive performance. The presence of cortical compensatory changes at prodromal stages of the disease is of high relevance, as they could contribute to the reduced effectiveness of therapeutic interventions^85^ that focus solely on the removal of pathological depositions.

## Acknowledgements

We would like to thank members of the Kuchibhotla lab for feedback on the manuscript, S. Ostojic for computational assistance, and J. Garmon for technical assistance. This work was supported by an Alzheimer’s Association Postdoctoral Fellowship (AS), a Kavli NDI Distinguished Postdoctoral Fellowship (JL), the HopkinsPREP NIH Training program (R25 GM109441) (KAF), a Provost’s Undergraduate Research Award (AW), an Albstein Research Scholarship (JA), David M. Rubenstein Fund for Hearing Research (AML), T32DC000023 (AML), R03AG081747 (AML), American Federation of Aging Research (KVK), and R01DC018650-02S1 (KVK).

## Author contributions

KVK, AS, and KAF designed the experimental approach. AS, SM, JL, KAF, AW, AML, KB and JA performed experiments. SM performed computational modeling. AS, SM analyzed behavioral data. AS, AW analyzed neural data. AS and KVK wrote the manuscript and all authors provided feedback.

## Methods

### Animals

All procedures were approved by Johns Hopkins University Animal Care and Use Committee. Male and female heterozygous mice (2 to 3-months old, n=6; 6 to 8-months-old, n=13; 10 to 12-months old, n=10) of a double transgenic mice that express chimeric mouse/human amyloid precursor protein and a mutant presenilin 1 (Jax B6;C3-Tg(APPswe,PSEN1dE9) 85Dbo/Mmjax, strain #034829-JAX) and litter-mate controls (2 to 3-months old, n=6; 6 to 8- months-old, n=12; 10 to 12-months old, n=10) were used for the behavior experiments. A subset of the 6-8mo mice used for behavior were used for the imaging experiments (APP/PS1- n=3 and APP/PS1+ n=5), and a group of 11mo APP/PS1+ (n=5) and APP/PS1- (n=5) were used to assess peripheral hearing. Animals were bred in house from JAX^®^ breeding pairs and housed in groups of 2-5 mice per cage and kept in a reverse light/dark cycle (10:30 am / 10:30 pm) with controlled temperature (19.5-22◦C) and humidity (35-38%).

### ABR measurements

Auditory Brainstem Responses were used to evaluate overall subcortical auditory function. Procedure was similar to what has been previously described ^86–88^. Briefly, mice were anesthetized i.p. with 95mg/kg ketamine and 9.5mg/kg xylazine and placed on an electronically controlled heating pad (DC controller FHC). Temperature was monitored rectally and maintained at 36°±1°C. Eye ointment was applied. Mice were placed inside a custom-made sound-attenuating box 10cm from a speaker (MF1, Tucker-Davis Technologies), measured from the left pinnae. Recordings were obtained using disposable subdermal needle electrodes (Rochester) placed over the animal vertex (active electrode), the left bulla (reference electrode) and the ipsilateral leg (ground electrode). Responses were amplified with a low nowise amplifier (Medusa4Z, Tucker-Davis Technologies, 100x) and digitally processed (RZ6-A-P1, Tucker-Davis Technologies). Responses were acquired with a sampling rate of 25kHz and offline band pass filtered (HP 300Hz, LP 3kHz). Tone and click stimuli were programmed, delivered and synchronized by the same processor (RZ6-A-P1, Tucker-Davis Technologies) using the *SigGenRZ* and *BioSigRZ* softwares (Tucker-Davis Technologies). Stimuli were presented at a rate of 21 repetitions/s, in 10dB decreasing intensities (from 90dB to 10dB) and presented 512 times. Click stimuli consisted of 0.1ms square wave pulses of alternating polarity. Tones (4, 8, 12, 16, 24, 32, 38, 42kHz) consisted of 5ms pulses (0.5ms on/off ramp).

### Surgical procedures

Mice were anesthetized with isoflurane (5.0% induction, 1.5-2.0% maintenance) and placed on a stereotactic apparatus (Kopf). Core body temperature was kept at 36°±1°C throughout the surgery. Eye ointment was applied and an antiseptic was used to clean the skin. After skull exposure, all connective tissue was removed with 3% hydrogen peroxide. For the behavioral experiments, a custom-made stainless steel headpost was attached to the skull with C&B Metabond dental cement (Parkell). For the imaging experiments, a 3mm craniotomy was centered 1.75mm anterior to lambda on the left ridge line. Then, 1μL of recombinant adeno-associated virus (dilution 1:15) encoding the calcium indicator GCaMP6f under the CamKII promoter (pENN.AAV.CamKII.GCaMP6f.WPRE.SV40, from James M. Wilson, Addgene, #100834-AAV9; http://n2t.net/addgene:100834; RRID:Addgene_100834; titer ≥ 1×10^13^ vg/mL) was injected into layer 2/3 of the left primary auditory cortex at a rate of 0.75μL/min with a 34G needle (1 inch, 12 degree bevel) and a 5μL capacity Hamilton syringe and a microinjection pump (Harvard Apparatus). After the virus was delivered, the needle was left in place for 8 min before removing it to allow diffusion of the viral particles. The exposed brain area was covered with a circular 3mm diameter glass window (Warner Instruments) and glued to the skull with *Krazy Glue*. Once dry, a custom-made stainless steel headpost was attached to the skull with C&B *Metabond* dental cement (Parkell). Subdermal Buprenorphine Base (1.0 mg/kg; Extended-Release Polymer Injection; Wedgewood) was administered for post-surgery analgesia.

### Auditory Go/No-Go discrimination task

After recovery from surgery, mice were progressively habituated to handling and head fixation for a total of 10 days. Mice were water-restricted and weighed daily. Each mouse received ∼1mL a day in order to maintain 80-85% of their original weight. Once habituated, mice underwent 2 days of lick training where they were placed in a plexiglass tube with an attached custom-made head fixation apparatus and trained to lick from a lick tube placed in front them without any stimulus presentation. Each lick was rewarded with 3uL of tap water. Mice were allowed to lick for 45 min or until they had consumed 1mL. On the next session, mice immediately began training on an auditory go/no-go task. Custom MATLAB (MathWorks) scripts were used to monitor all behavioral events and to interface with BPOD State Machines (r1 or r2, Sanworks) and control stimulus presentation and reward delivery. Licks were detected with a custom-made infrared photogate. A free field electrostatic speaker (ES1, Tucker-Davis Technologies) was located ∼5 cm from the animal’s left ear and was driven by an electrostatic speaker driver (E1, Tucker-Davis Technologies). S+ and S- tones used were one quarter octave apart and range from 4757 to 38000 Hz. Tones were calibrated to an intensity of 65-70 dB (SPL). The head fixation apparatus and speaker were enclosed in a custom-made sound-attenuated box. Each animal received 6-7 training sessions per week. Each session consisted of the presentation of 320 trials where S+ and S- tones were pseudo-randomly ordered every 20 trials so both tones were played on equal number of trials per session. Each trial had a pre-stimulus no-lick period (2s) that was followed by the stimulus presentation (100ms), a delay (100ms), a response period (2s) and variable inter-trial interval depending on trial outcome (Fig**. 1b**). S+ trials where the animal licked were rewarded with 3uL of water. Every other day, for 20 of the total trials in the middle of the session, the lick tube was retracted and reward was not delivered (probe trials). Training continued until mice reached plateau performance for 3 days.

### Two-photon calcium imaging

After 2-3 weeks of viral expression and water restriction we started the imaging experiments. Mice were habituated and head-fixed in a clear plexiglass tube within a sound attenuated chamber. On the first day of imaging, a pseudo-random sequence of 17 pure tones (4-to-64 kHz; duration 100ms) spaced one quarter octave were played 10 times at 70dB through an electrostatic speaker driver (RZ6, TDT) to a free field electrostatic speaker (ES1, Tucker-Davis Technologies). Tones were selected for behavioral training following two conditions, that none of them were the BF of the field of view nor the same frequency as the galvo-resonant scanner (8kHz). On consecutive days mice started behavioral training as described in the auditory go/no-go discrimination task section. Two-photon resonant-scanning microscope (Neurolabware) was used for imaging. GCamp6f (calcium sensor) and Methoxy-X04 (β-amyloid dye) were excited at 980nm and 860nm respectively using an Insight X3 laser (Spectra-Physics) with emission collected using green and blue channels. Images were collected using ScanBox (Neurolabware) with a rotatable objective (16x, 0.8NA, Nikon) set at ∼50 degrees to image the auditory cortex. For 3-D plaque imaging, a z-stack of 400μm was recorded at 1x by imaging 50 frames every 2μm. For functional imaging, 2 or 4 planes 70μm apart and ∼200μm below the dura (layer 2/3) were imaged using an electronically tunable lens in the center of the larger plaque volume and imaged at 2x (0.796mm X 0.512mm). Data was collected at 31.25 frames per second. Methoxy X04 (10mg/kg) was administered i.p. 24 hours before every imaging session.

### Data analysis

Data processing and statistical analysis was performed using custom scripts written in MATLAB (MathWorks) or GraphPad Prism. Data was tested for normality and parametric or non-parametric statistical tests were applied accordingly as described in the figure legends. A mixed-effects model was used for repeated measures data with unequal number of data points. Plots show mean ± s.e.m.

#### Behavior

Hits and false alarm action rates were measured in blocks 20 trials for comparison of reinforced and probe blocks. Performance (*d’*) was calculated by subtracting the z-scored false alarm rate to the z-scored hit rate. Action rates were corrected by 1-1/2N or 1/2N when 1 or 0 respectively to avoid infinite values. Reaction time was defined as the time of the first lick after tone presentation.

#### ABRs

Threshold detection for all stimulus and mice was performed by 4 people blind to mice genotypes and was defined as the last dB SPL at which any wave was detected. The peak and valley of the first 5 waves of the ABR trace were manually selected using *BioSigRZ* software (Tucker-Davis Technologies). A mixed-effects model was used to test for significance accounting for the missing latencies and amplitudes due to the absence of responses to some intensities or frequencies.

#### Two-photon imaging

Motion correction and non-rigid registration was performed with *suite2p* (https://github.com/MouseLand/suite2p). Neural traces were obtained from *suite2p-cellpose*- detected regions of interest (ROI). Change in fluorescent trace was measured as, ΔF =(F-F0)/F0, with F being the mean fluoresce during the response window and F0 the mean activity during 1s of baseline before the tone presentation. ROIs were considered responsive when their evoked activity was significantly different across all presentations of the same stimulus/trial type (p<0.01, sided Wilcoxon signed rank test). The selectivity Index was calculated by dividing the average response of each neuron to the S^+^ minus the average response of the same cell to the S^-^ by the sum of the absolute activity in response to both. Peak detection was performed with MATLAB function *findpeaks* on smoothed traces. For reinforced versus probe comparisons, only neurons of mice with probe performance (d’) higher than 1.2 were selected for the analysis. A linear discriminant classifier was used as described ^64^ to decode the stimulus presented. Population vectors CT,t and CF,t were calculated from a random selection consisting of ½ of the total S^+^ and S^-^ trials. If the total trials were not an even number, then the number was rounded to the nearest integer less than that number. Each time bin was defined as one frame and a decoding vector Wt was obtained as follows,

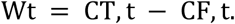

This was used together with the bias

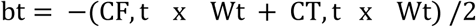

as a decision rule for the test population activity vectors obtained from an equal number of the remaining S^+^ and S^-^ test trials that were not used to train the classifier.

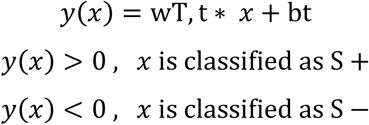

#### Distance to plaque analysis

An image segmentation of plaques from each z-stack plane was obtained with *Ilastik* ^89^ and FIJI ^90^. We performed a 3D distance transformation to assign each non-plaque pixel with a value that equaled the distance to the nearest non-zero value (i.e. the nearest plaque). Neuron coordinates were extracted from *suite2p*. The mean image of the calcium signal from each behavioral session was registered to the session z-stack and the centroid of each ROI was used to calculate the distance to the closest plaque.

#### Reinforcement learning model

We adapted a previously constructed reinforcement learning model that accounts for contextual modulation ^22^. Briefly, the model is governed by 9 parameters. A sensory coding population (Extended Data Fig. 7a) that is selective to the S^+^ (green) or S^-^ (orange) tones, or non-selective (S, black). Sensory outputs converge in one inhibitory (I) and one excitatory (D) decision making population that is modulated by a contextual scaling factor (yellow) in the reinforced condition only. Reward signals, present only in one condition, modify the weights of the sensory population to favor the correct answer. Two noise factors scale the decision-making population (σ) or the weights (k). S^+^ and S^-^ learning rates (αr αnr) also modify the weights of the decision-making populations.

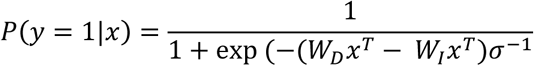

We ran 2000 model simulations using a Bayesian adaptive search (BADS) using the action rates for each individual animal (n= 12 APP/PS1-, n= 13 APP/PS1+, 6-8 months old). The best 50 simulations were chosen for each individual animal using the goodness of fit. We plotted the distribution of the 9 model-parameters for control and APP/PS1^+^ mice, and only those showing a statistically different distribution (Chi squared test) were considered as the parameters governing the differences between groups.

---

**Extended Data Figure 1.**
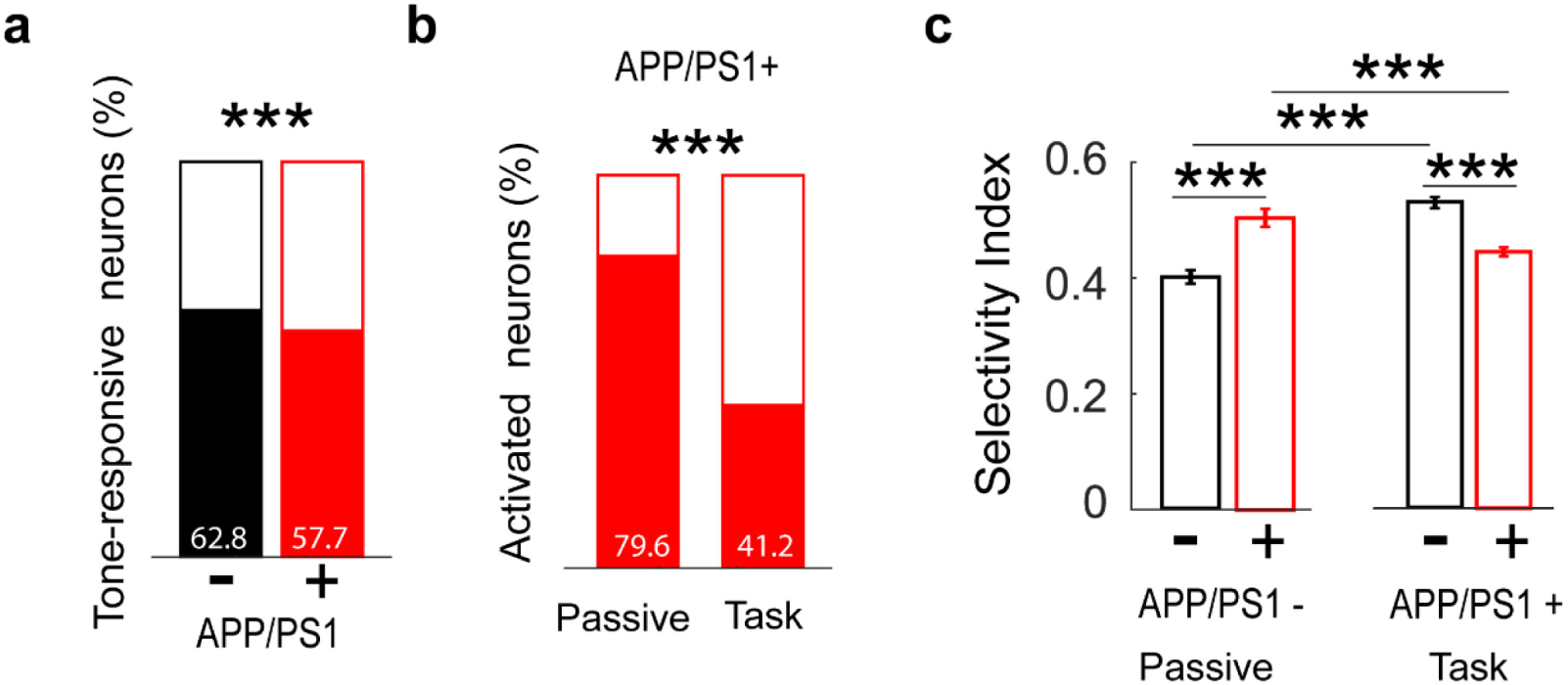
**a,** Tone responsiveness before training is slightly impaired (5.1% decrease) in APP/PS1+ mice compared to controls (χ2=18.3576; p<0.001). **b,** While the majority of responsive neurons is activated both in the passive and during the task for control mice, this only occurs in the passive context for APP/PS1+ mice (χ2=221.8577; p<0.00001). **c,** Selectivity of significantly responsive neurons is higher in APP/PS1+ neurons compared to controls before training (passive) but lower than control after training (control vs APP/PS1+ in passive Z=-5.7146, p<0.001; control vs APP/PS1+ task Z =8.1777, p=2.8931e- 16; APP/PS1- before and after training: Z=-8.6376; p<0.001; APP/PS1+ before and after training (Z=4.4655; p<0.001)). ∗ p<0.05, ∗∗ p<0.01, ∗∗∗ p<0.001, ns=non-significant.

**Extended Data Figure 2.**
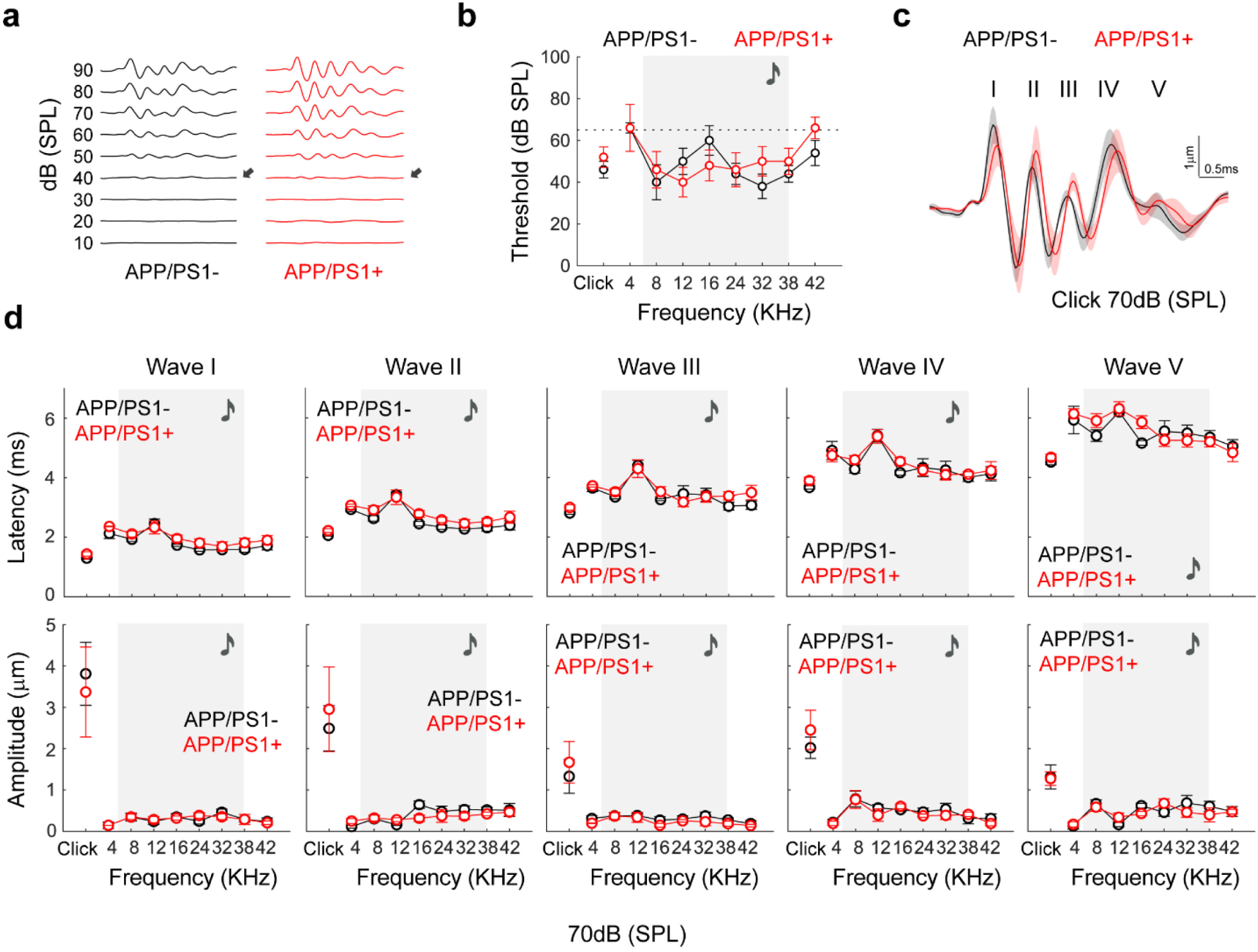
APP/PS1+ mice subcortical and peripheral hearing is preserved. **a,** Exemplar auditory brainstem responses to clicks. Arrows indicate threshold. **b,** Auditory detection thresholds to clicks and tones. Shading indicates tone frequencies used in the auditory go/no go task (2- way ANOVA; Interaction, F(8,64)=1.551, p=0.16; Stimulus, F(3.429,27.43)=5.425, p=0.0034; Genotype, F(1,8)=0.1264, p=0.7314). **c,** Grand average across all mice of the ABR trace in response to a click of 70dB SPL. **d,** Latencies and amplitudes of the 5 first peaks to every stimulus at 70db SPL (11-month-old mice, n=5 APP/PS1-, n=5 APP/PS1+, see stats in table S1. ∗ p<0.05, ∗∗ p<0.01, ∗∗∗ p<0.001, ns=non-significant.

**Extended Data Figure 3.**
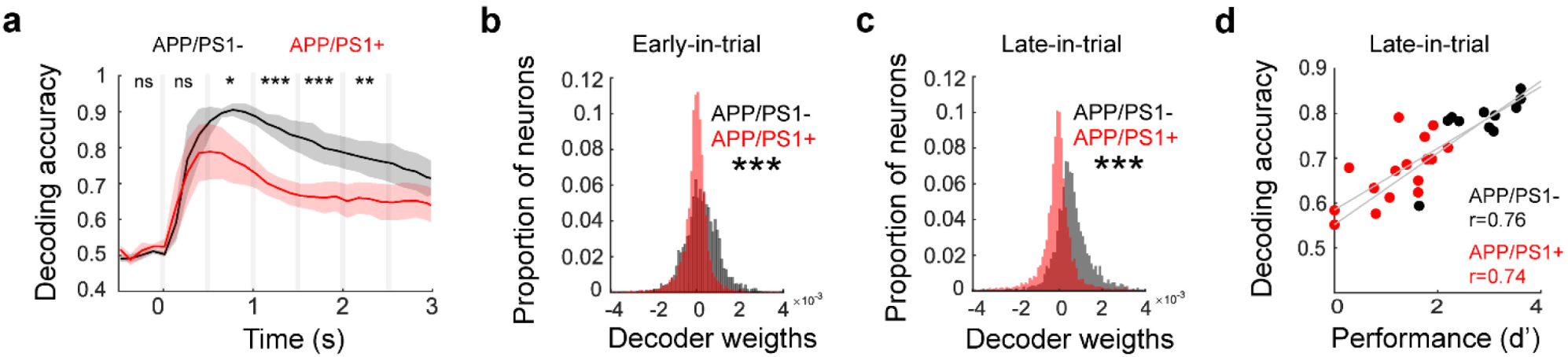
Tone decoding impairments in 6-8mo APP/PS1+ mice. **a,** Tone decoding is partially preserved early-in trial but worsens late-in-trial (0.5-1s Genotype, F(1,24)=7.155, p=0.0132; 1-1.5s Genotype, F(1,24)=39.66, p<0.0001; 1.5-2s Genotype, F(1,24)=22.40, p<0.0001; 2-2.5 Genotype, F(1,24)=11.22, p=0.002; n=3 APP/PS1-, n=5 APP/PS1+). **b-c,** Distribution of decoding weights early and late-in-trial show fewer neurons that contribute to the stimulus decoding in APP/PS1+ mice (p<0.001 early-in-trial; p<0.001 late-in-trial; Two-sample Kolmogorov-Smirnov; n=2,173 APP/PS1-, n=4,043 APP/PS1+). **d**, Correlation between decoding accuracy in 100-trials block and task performance. APP/PS1- p=0.0063; APP/PS1+ p=0.0007.∗ p<0.05, ∗∗ p<0.01, ∗∗∗ p<0.001, ns=non-significant.

**Extended Data Figure 4.**
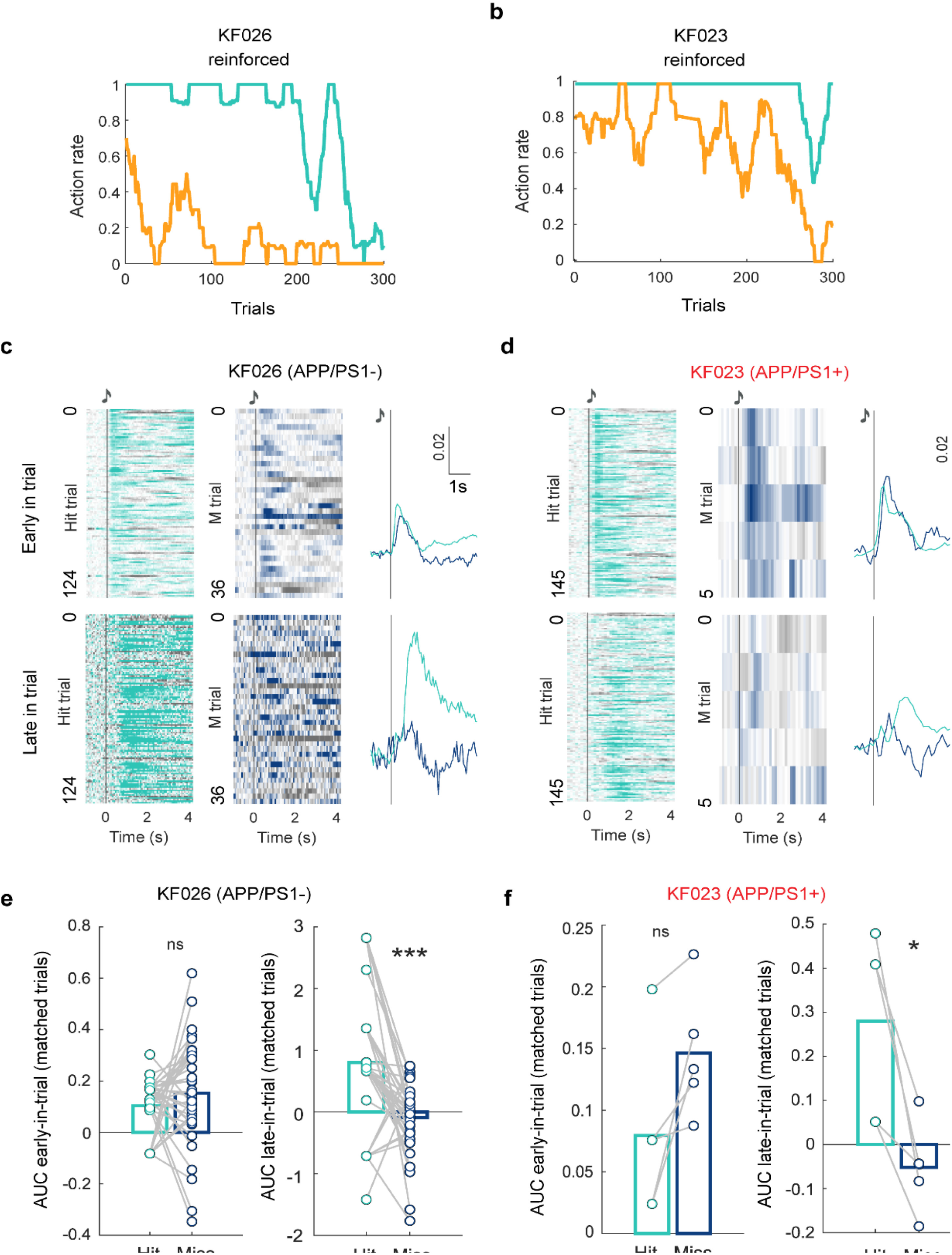
Late-in-trial activity is not a pure sensory response. **a,** Behavior during imaging of a 6-8mo control mouse in **c** and **e**. **b,** Behavior during imaging of an APP/PS1+ mouse in **d** and **f**. **c,** Heatmap of neural responses to all hit and all miss trials of significantly activated neurons early-in-trial (n=639) and late-in-trial (n=11) of a control example mouse. **d,** Heatmap of neural responses to all hit and all miss trials of significantly activated neurons early-in-trial (n=202) and late-in-trial (n=94) of an APP/PS1+ example mouse. **e,** Area under the curve of all significant neurons of early-in-trial activity for every hit trial immediately before a miss and for every miss trial in the example control mouse showed in **a** and **c** (early-in-trial, p=0.27145; late-in-trial, p=0.000615); **f,** Area under the curve of all significant neurons of early-in-trial activity for every hit trial immediately before a miss and for every miss trial in the APP/PS1+ mouse showed in **b** and **d** (early-in-trial, p=0.051332; late-in-trial, p=0.015). ∗ p<0.05, ∗∗ p<0.01, ∗∗∗ p<0.001, ns=non- significant.

**Extended Data Figure 5.**
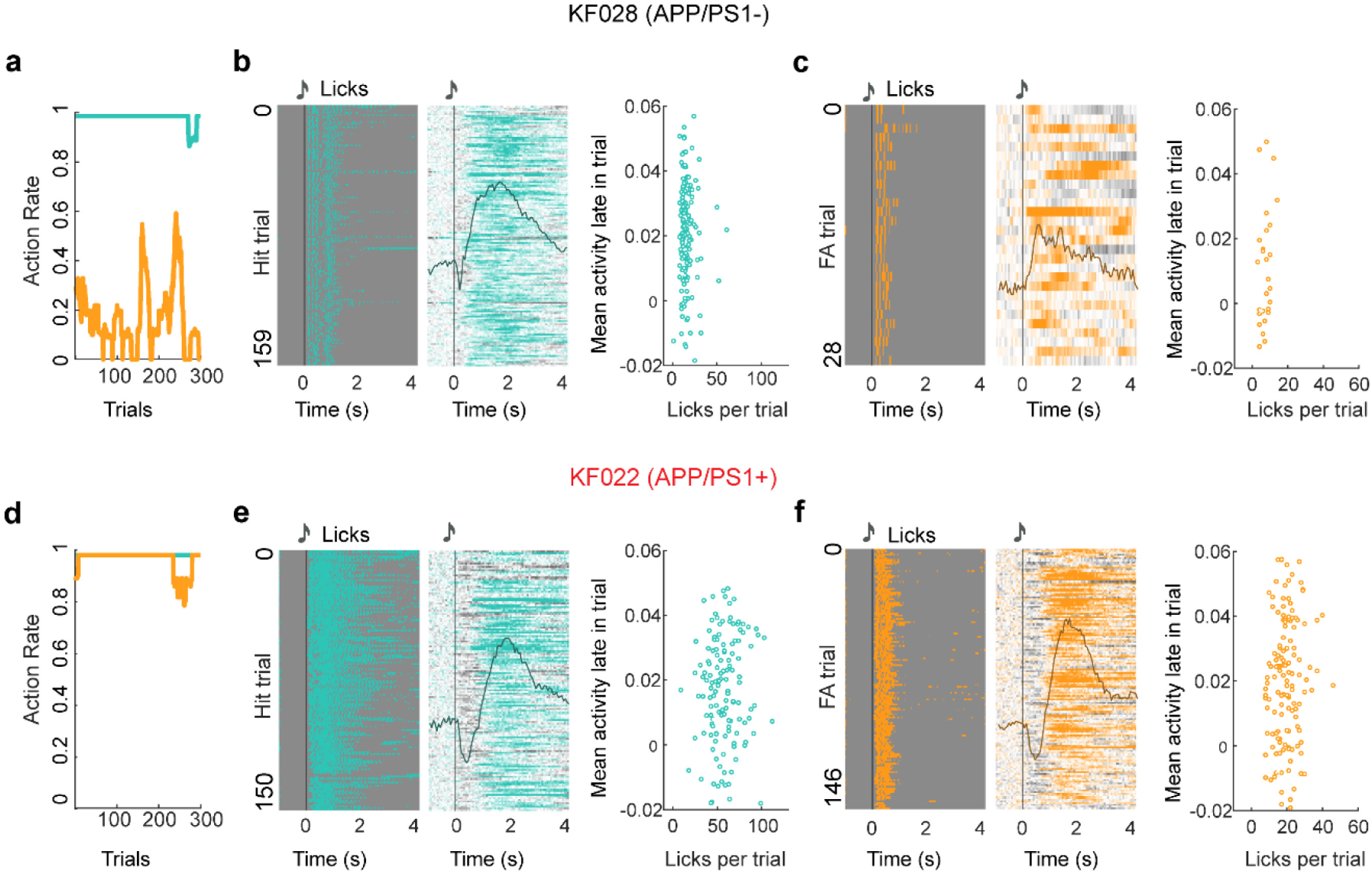
Activity late-in-trial is independent of lick vigor within trial. **a,** Behavior during imaging in exemplar control mouse. **b,** Licks for every hit trial in A. **c,** Heatmap and overlay average of neural responses of putative reward prediction neurons late-in-trial per hit trial (n=139 neurons). **d,** Mean neural activity in every hit trial during the late-in-trial period (y-axis) is independent of number of licks of that trial (x-axis) (r=0.02; p=0.8023). **e,** Licks for every FA trial in A. **f,** Heatmap and overlay average of neural responses of putative reward prediction neurons late-in-trial per FA trial (n=139 neurons). **g,** Mean neural activity in every false alarm trial during the late-in-trial period (y-axis) is independent of number of licks of that trial (x-axis) (r=0.3292; p=0.0872). **h,** Behavior during imaging in exemplar APP/PS+ mouse. **i,** Licks for every hit trial in H. **j,** Heatmap and overlay average of neural responses of reward prediction neurons late-in-trial per hit trial (n=50 neurons). **k,** Mean neural activity in every hit trial during the late-in-trial period (y-axis) is independent of number of licks of that trial (x-axis) (r=0.0917; p=0.2711). **l,** Licks for every FA trial in H. **m,** Heatmap and overlay average of neural responses of reward prediction neurons late-in-trial per FA trial. **n,** Mean neural activity in every false alarm trial during the late-in-trial period (y-axis) is independent of number of licks of that trial (x-axis) (r=0.0647; p=0.9353). ∗ p<0.05, ∗∗ p<0.01, ∗∗∗ p<0.001, ns=non-significant.

**Extended Data Figure 6.**
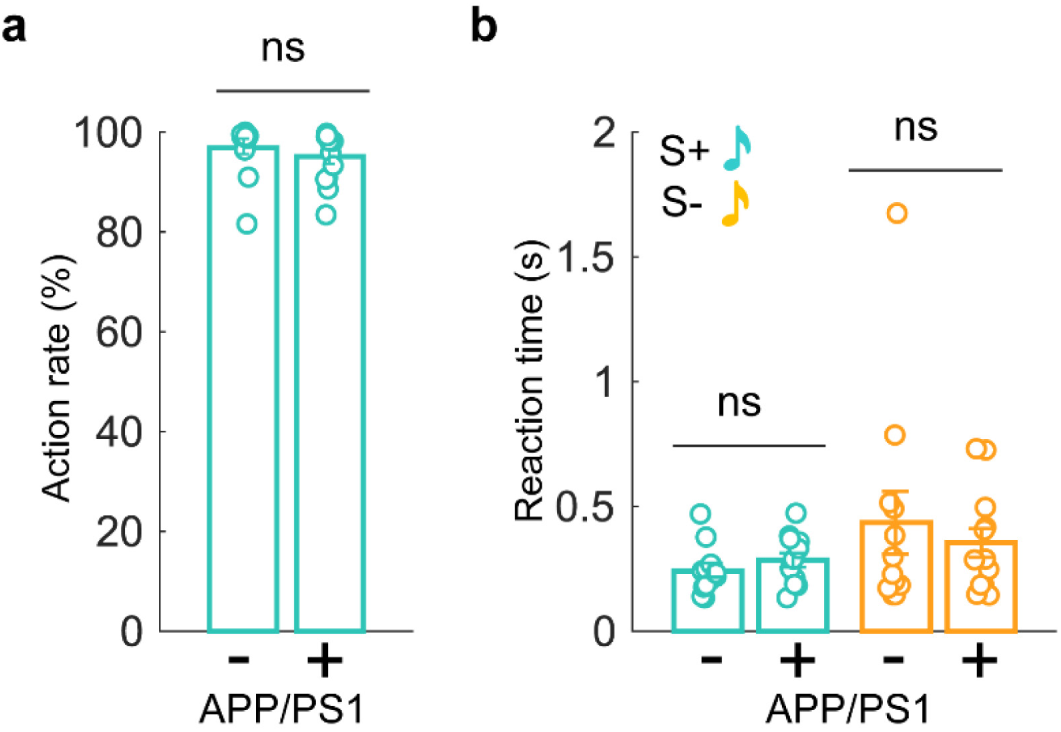
6-8mo amnestic mice show no evidence of disengagement, impulsivity or motor-related licking ability. **a,** Action rate on S+ trials is not significantly different between APP/PS1+ mice and age matched littermates (Z=0.8975, p=0.369, Wilcoxon rank-sum test). **b,** APP/PS1+ mice have similar reaction times compared to control mice (Z=-1.241, p=0.215 hits; Z=0, p=1 false alarms). n=12 (APP/PS1-, 6-8mo), n=13 (APP/PS1+, 6-8mo); ∗ p<0.05, ∗∗ p<0.01, ∗∗∗ p<0.001, ns=non-significant.

**Extended Data Figure 7.**
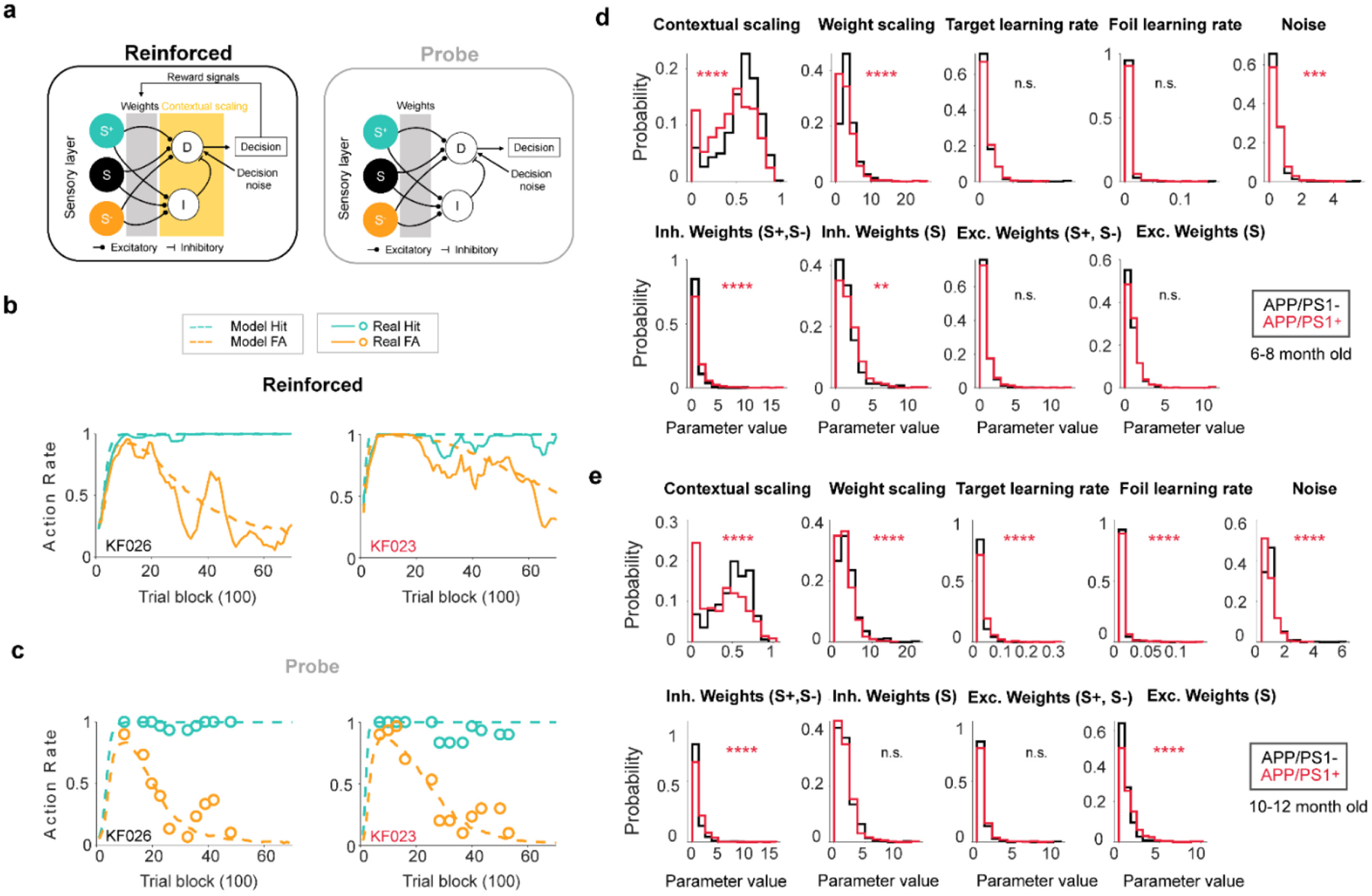
Contextual scaling and inhibitory weights are the main drivers of performance deficits on reinforced trials in amnestic mice. **a,** Schematic of the model. The model is governed by 9 parameters. A sensory coding population that is selective to the S+ (green) or S-(orange) tones, or non-selective (S, black). Sensory outputs converge in one inhibitory (I) and one excitatory (D) decision making population that is modulated by a contextual scaling factor (yellow) in the reinforced condition only. Reward signals, present only in the reinforced condition, modify the weights of the sensory population to favor the correct answer. Two noise factors scale the decision-making population (σ) or the weights (k). S+ and S- learning rates (αr αnr) also modify the weights of the decision-making populations. **b,** Model fit of the action rates in a control and APP/PS1+ mice (6 to 8-months-old) in the reinforced context. 100 trials per trial block. **c**, Model fit of the action rates in a control and APP/PS1+ mice (6 to 8-months-old) in the probe context. 100 trials per trial block. **d,** Contextual scaling and inhibitory neuron parameters are most impacted in APP/PS1+ mice; Contextual scaling, p=1.16e-14; Weight scaling, p=1.11e-9; S+ learning rate, p=0.16; S- learning rate, p=0.37; Noise, p=0.00071; Inhibitory weights of the S+ and S- populations, p=1.81e-5; Inhibitory weights of the S population, p=0.0012; Excitatory weights of the S+ and S- populations, p=0.33; Excitatory weights of the S population, p=0.082; Wilcoxon rank sum tests; n=12 APP/PS1-, n=13 APP/PS1+. **e,** Parameters impacted in 10-12mo APP/PS1+ mice; Contextual scaling, p=5.97e-15; Weight scaling, p=8.4e-6; S+ learning rate, p=1.06e-5; S- learning rate, p=2.86e-8; Noise, p=1.18e-8; Inhibitory weights of the S+ and S- populations, p=9.36e-8; Inhibitory weights of the S population, p=0.68; Excitatory weights of the S+ and S- populations, p=0.41; Excitatory weights of the S population, p=1.11e-5; n=10 APP/PS1-, n=11 APP/PS1+. ∗ p<0.05, ∗∗ p<0.01, ∗∗∗ p<0.001, ns=non-significant.

**Extended Data Figure 8.**
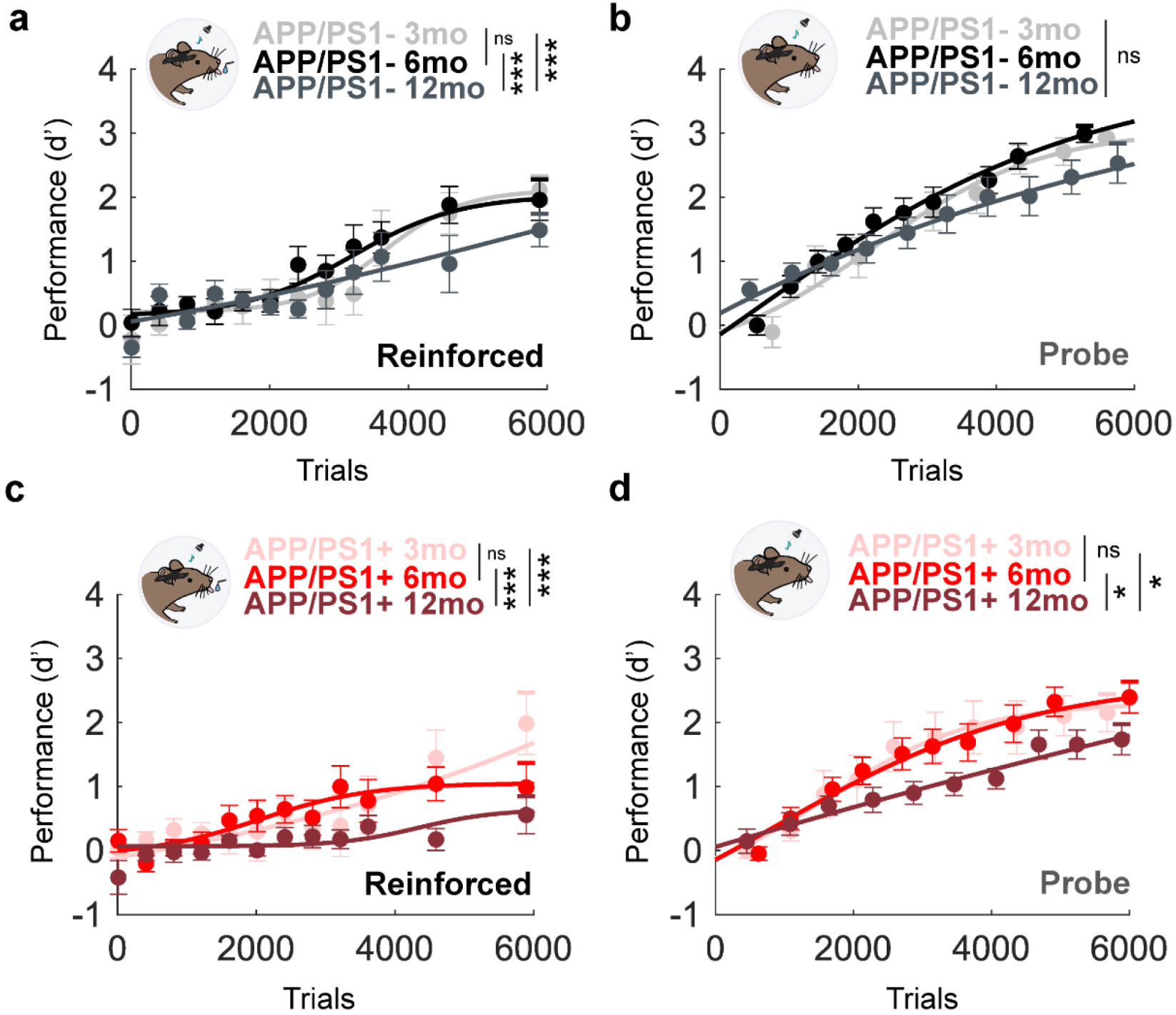
Deficits in contextual expression are age-dependent and decline faster in amnestic mice. **a,** Control mice performance in the reinforced context (Mixed-effects model; Interaction, F(118,1456)=1.539, p=0.0003; 2-3 vs 6-8mo, p=0.7894; 2-3 vs 10-12mo, p<0.0001; 6-8 vs 10-12mo, p<0.0001). **b,** Sensorimotor memories at 2-3, 6-8 and 10-12 months-old in control mice (Mixed-effects model; Interaction, F(24,278)=2.349, p=0.0005; 2-3 vs 6-8mo, p=0.5436; 2-3 vs 10-12mo, p=0.151; 6-8 vs 10- 12mo, p=0.5163). **c,** Performance deficits in APP/PS1^+^ mice (Mixed-effects model; Interaction, F(118,1555) =1.967, p<0.0001; 2-3 vs 6-8mo, p=0.9523; 2-3 vs 10-12mo, p<0.0001; 6-8 vs 10-12mo, p<0.0001). **d,** Sensorimotor memories are preserved in APP/PS1+ mice but degrade at 10-12mo (Mixed-effects model; Interaction, F(12,162)=1.837, p=0.0463; 2-3 vs 6-8mo, p=0.9961; 2-3 vs 10-12mo, p=0.1169; 6-8 vs 12mo, p=0.0217). APP/PS1-: n=6 (2-3mo); n=12 (6-8mo); n=10 (10-12mo). APP/PS1+ mice: n=6 (2-3mo); n=13 (6-8mo); n=11 (10-12mo). ∗ p<0.05, ∗∗ p<0.01, ∗∗∗ p<0.001, ns=non-significant).

**Table 1.**
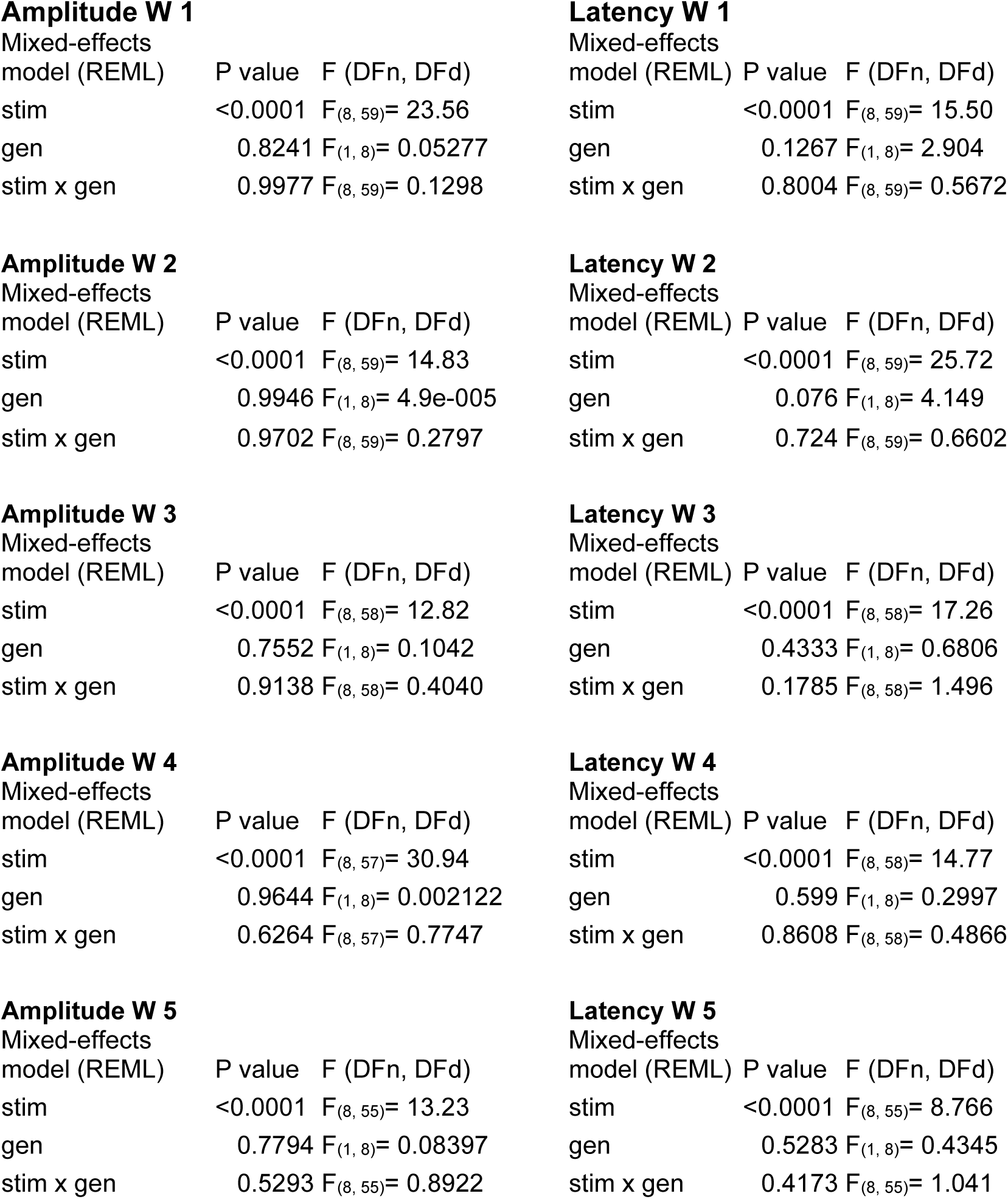
Statistics for Extended Data Figure 2D.

**Amplitude W 1**
| Mixed-effects<br>model (REML) | P value | F (DFn, DFd) |
| --- | --- | --- |
| stim | <0.0001 | $F_{(8, 59)} = 23.56$ |
| gen | 0.8241 | $F_{(1, 8)} = 0.05277$ |
| stim x gen | 0.9977 | $F_{(8, 59)} = 0.1298$ |

**Amplitude W 2**
| Mixed-effects<br>model (REML) | P value | F (DFn, DFd) |
| --- | --- | --- |
| stim | <0.0001 | $F_{(8, 59)} = 14.83$ |
| gen | 0.9946 | $F_{(1, 8)} = 4.9\text{e-}005$ |
| stim x gen | 0.9702 | $F_{(8, 59)} = 0.2797$ |

**Amplitude W 3**
| Mixed-effects<br>model (REML) | P value | F (DFn, DFd) |
| --- | --- | --- |
| stim | <0.0001 | $F_{(8, 58)} = 12.82$ |
| gen | 0.7552 | $F_{(1, 8)} = 0.1042$ |
| stim x gen | 0.9138 | $F_{(8, 58)} = 0.4040$ |

**Amplitude W 4**
| Mixed-effects<br>model (REML) | P value | F (DFn, DFd) |
| --- | --- | --- |
| stim | <0.0001 | $F_{(8, 57)} = 30.94$ |
| gen | 0.9644 | $F_{(1, 8)} = 0.002122$ |
| stim x gen | 0.6264 | $F_{(8, 57)} = 0.7747$ |

**Amplitude W 5**
| Mixed-effects<br>model (REML) | P value | F (DFn, DFd) |
| --- | --- | --- |
| stim | <0.0001 | $F_{(8, 55)} = 13.23$ |
| gen | 0.7794 | $F_{(1, 8)} = 0.08397$ |
| stim x gen | 0.5293 | $F_{(8, 55)} = 0.8922$ |

**Latency W 1**
| Mixed-effects<br>model (REML) | P value | F (DFn, DFd) |
| --- | --- | --- |
| stim | <0.0001 | $F_{(8, 59)} = 15.50$ |
| gen | 0.1267 | $F_{(1, 8)} = 2.904$ |
| stim x gen | 0.8004 | $F_{(8, 59)} = 0.5672$ |

**Latency W 2**
| Mixed-effects<br>model (REML) | P value | F (DFn, DFd) |
| --- | --- | --- |
| stim | <0.0001 | $F_{(8, 59)} = 25.72$ |
| gen | 0.076 | $F_{(1, 8)} = 4.149$ |
| stim x gen | 0.724 | $F_{(8, 59)} = 0.6602$ |

**Latency W 3**
| Mixed-effects<br>model (REML) | P value | F (DFn, DFd) |
| --- | --- | --- |
| stim | <0.0001 | $F_{(8, 58)} = 17.26$ |
| gen | 0.4333 | $F_{(1, 8)} = 0.6806$ |
| stim x gen | 0.1785 | $F_{(8, 58)} = 1.496$ |

**Latency W 4**
| Mixed-effects<br>model (REML) | P value | F (DFn, DFd) |
| --- | --- | --- |
| stim | <0.0001 | $F_{(8, 58)} = 14.77$ |
| gen | 0.599 | $F_{(1, 8)} = 0.2997$ |
| stim x gen | 0.8608 | $F_{(8, 58)} = 0.4866$ |

**Table 1. Statistics for Extended Data Figure 2D**
| Mixed-effects<br>model (REML) | P value | F (DFn, DFd) |
| --- | --- | --- |
| stim | <0.0001 | $F_{(8, 55)} = 8.766$ |
| gen | 0.5283 | $F_{(1, 8)} = 0.4345$ |
| stim x gen | 0.4173 | $F_{(8, 55)} = 1.041$ |

**Table 2.** Probe performance. APP/PS1-, 6-8mo, n=12. APP/PS1+, 6-8mo, n=13.

|  | P value | F (DFn, DFd) |
| --- | --- | --- |
| Time | <0.0001 | $F_{(3,492, 76,01)} = 77.27$ |
| Genotype | 0.266 | $F_{(1, 23)} = 1.302$ |
| Interaction | 0.5 | $F_{(13, 283)} = 0.9512$ |

